# Unveiling abundance-dependent metabolic phenotypes of microbial communities

**DOI:** 10.1101/2023.01.10.523517

**Authors:** Natalia E. Jiménez, Vicente Acuña, María Paz Cortés, Damien Eveillard, Alejandro Maass

**Affiliations:** Centerfor Mathematical Modeling, University of Chile and CNRS-IRL2807, Santiago, Chile; Millennium Institute Centerfor Genome Regulation, University of Chile, Santiago, Chile; Department of Mathematical Engineering, University of Chile, Santiago, Chile; Nantes Université, Ecole Centrale Nantes, CNRS, LS2N, France

**Keywords:** constrained-based modeling, microbial communities, metabolic plasticity

## Abstract

Constraint-based modeling has risen as an alternative for characterizing the metabolism of communities. Adaptations of Flux Balance Analysis have been proposed to model metabolic interactions in most cases, considering a unique optimal flux distribution derived from the maximization of biomass production. However, these approaches do not consider the development of other potentially novel essential functions not directly related to cell growth which forces them to display suboptimal growth rates in nature. Additionally, suboptimal states allow a degree of *plasticity* in the metabolism, thus allowing quick shifts between alternative flux distributions as an initial response to environmental changes.

In this work, we present a method to explore the *abundance-growth space* as a representation of metabolic flux distributions of a community. This space is defined by the composition of a community, represented by its members’ relative abundance and their growth rate. The analysis of this space allows us to represent the whole set of feasible fluxes without needing a complete description of the solution space unveiling abundance-dependent metabolic phenotypes displayed in a given environment. As an illustration, we consider a community composed of two bioleaching bacteria, *Acidithiobacillus ferrooxidans* Wenelen and *Sulfobacillus thermosulfidooxidans* Cutipay, finding that changes in the composition of their available resources significantly affects their metabolic plasticity.

**IMPORTANCE:** In nature, organisms live in communities and not as isolated species. Their interactions provide a source of resilience to environmental disturbances. Despite their importance in ecology, human health, and industry, understanding how organisms interact in different environments remains an open question.

In this work, we provide a novel approach which, only using genomics information, studies the metabolic phenotype exhibited by communities, where the exploration of suboptimal growth flux distributions and the composition of a community allows to unveil its capacity to respond to environmental changes, shedding the light of the degree of metabolic plasticity inherent to the community.

## INTRODUCTION

In nature, organisms live in communities rather than as isolated species. These communities emerge from interactions (1) and provide a source of resilience to perturbations such as biofilm development (2, 3), cross-feeding (4) or community resistance to environmental disturbances (5). The study of these interactions is imperative in critical areas such as ecology (6), health (7), and the biotechnological industry (8). Understanding how organisms interact in different environments is still an open question despite their relevance.

The last decade saw the rise of metabolic modeling for formalizing microbial interactions. The metabolic cross-talking between microorganisms notably justifies the metabolic abstraction for sustaining biogeochemical cycles (9). As a natural following, several approaches have been developed for modeling the metabolism of communities (10) using comprehensive genome-scale descriptions for which each organism is described by its inner biochemical machinery (11, 12, 13, 14). This genome-scale description allows to develop several computational strategies to identify essential metabolic interactions (e.g., SteadyCom (15), RedCom (13), community Flux Balance Analysis (cFBA) (16), and MICOM (17)). When applied in specific contexts, it results in insightful advances like the determination of essential interactions in anaerobic digestion for biogas production (18), critical reactions associated with cancer (19), and the gut microbiota (20).

These approaches characterize metabolic flux distributions, and community compositions using an adaptation of Flux Balance Analysis (FBA) (21), where an objective function, usually biomass production, is optimized (22, 15, 13, 23, 16, 17). Maximization of biomass is of great interest for growth rate estimation, and it has shown substantial benefits in the biotechnological context that aims at controlling single strains to improve yields for a product of interest (24, 25). However, such a modeling assumption does not stand in a natural environment. In the case of microbial communities, biomass maximization does not always account for secondary metabolites essential to sustain communities via metabolic cross-talking (26) and protect against environmental perturbations (5).

A challenge for studying and predicting metabolic interactions in community models is to propose methods allowing the exploration of alternative metabolic states that a community could display while preserving the practicality of traditional methods like FBA and Flux Variability Analysis (FVA (27)). A recent alternative approach was proposed to directly explore the flux space defined by exchange reactions to characterize its metabolic niche (28). Comparison between this newly defined space for different organisms allows for determining environments where a community can thrive. However, this approach still leaves out the role and abundance of each organism in the community. Indeed, the relative abundance of members of a community shape the distribution of resources and synthesis of secondary metabolites (15). Although there are promising ideas regarding the laws governing its probabilistic distribution (29), the prediction of abundances is uncertain at steady state. Thus exploration of the effect of community composition is critical for studying metabolic interactions.

In this work, we explore the space of metabolic fluxes of a community, focusing on the distribution of abundances of its organisms and community growth rates. This space is partitioned according to displayed metabolic phenotypes compatible with each abundance-growth point, computed based on flux ranges for each reaction. These ranges are seen as descriptors of the system’s plasticity (30, 31), and allow us to distinguish between reactions that present *flux plasticity*, where their fluxes remain positive (i.e., always active) despite flux variations, from reactions associated with *structural plasticity* that show zero flux on some solutions, emphasizing their putative replacement by alternative pathways.

This novel framework allows projecting the community metabolic fluxes in a lower dimensional space for interpretation, thus providing a practical exploration of metabolic phenotypes in a community characterized by different relative abundances and growing at suboptimal growth rates. For illustration, this method is applied to a bioleaching community composed by *Acidithiobacillus ferrooxidans* Wenelen and *Sulfobacillus thermosulfidooxidans* Cutipay, allowing to identify of predominant metabolic pathways activated in different zones of the community flux space.

## RESULTS

### Rationale of the community metabolic phenotype: the abundance-growth space

We aim to understand mechanisms that support community structure and the ecosystem’s resilience to perturbations. To achieve this, we explore metabolic phenotype changes displayed by a community of *N* organisms subjected to relative abundances and growth rate variations. A community model is constructed using single-organism genome-scale metabolic models, considering each organism as a compartment within the ecosystem. Exchanges between organisms occur in the pool compartment, and all organisms grow at the same growth rate to maintain constant relative abundances over time (16, 13, 15).

In this model, *μ* is the community growth rate, and the relative abundance *f_i_* of organism *i* appears as a factor of the flux bounds for any reaction of its respective organism (Methods). Thus, variations on the vector of abundances *F* = (*f*_1_,…,*f_N_*) and on *μ* affect the set of flux distributions that are feasible solutions of the model. Specifically, for a given abundance-growth pair (*F,μ*), the *polytope P_F,μ_* denotes all flux distributions satisfying the model’s constraints. If *P_F,μ_* is empty, no feasible flux distribution yields a growth rate μfor a vector of abundances F. On the other hand, if there is at least one flux distribution in *P_F,μ_*, the abundance-growth pair (*F,μ*) is *feasible*. Hence, the *abundance-growth space* is the set of all feasible pairs (*F,μ*).

To explore variations of phenotypes occurring for different abundances F and growth rates μ, we analyze how *P_F,μ_* varies across the abundance-growth space. A numerical description of this space is achieved by computing a grid 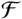 of feasible abundance-growth points (Methods). In the case of a community of two organisms, this grid is two dimensional (*f*_1_,*μ*) since *f*_2_ = 1 − *f*_1_ (Figure 1 (b)).

**FIG 1.**
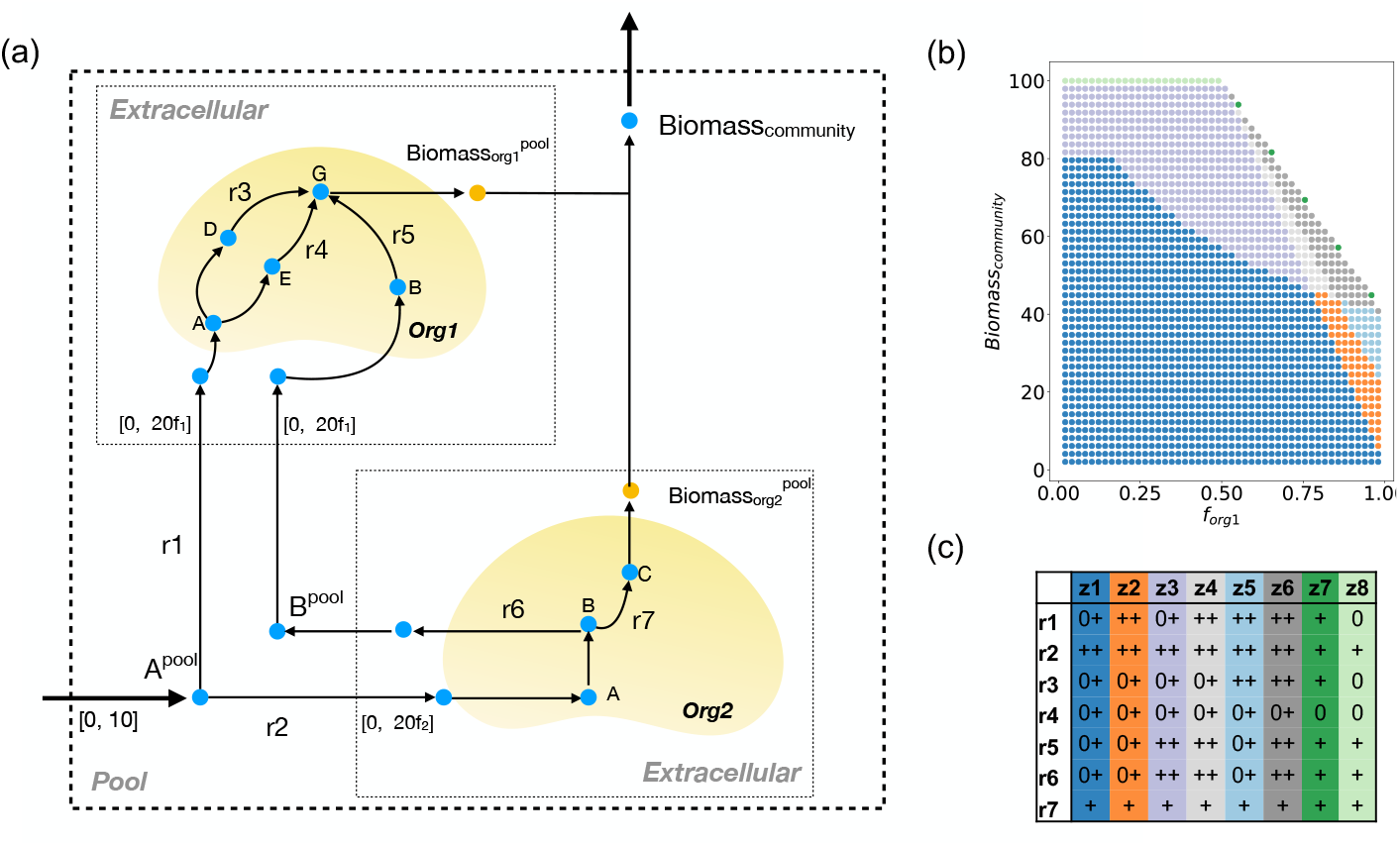
Illustration of a community metabolic phenotypic space. (a) One assumes a microbial community metabolic network composed of two organisms. The community model shows two extracellular compartments for each organism and a pool compartment and their exchange reactions. Note that substrate uptake is constrained by the composition of the community (Methods). By consuming metabolite A, org2 can grow or produce metabolite B, while org1 grows by consuming A or B. The set *R′* of selected reactions is composed by *r_i_* with *i* ∈ {1,…,7}. (b) Abundance-growth space and the partition associated with the set of selected reactions. (c) Categorical vectors associated with reactions on each zone. Qualitative changes are associated with increased biomass production and changes in community composition. For instance, structural plasticity (0+) of organism 1 in zone 1 is gradually replaced by flux plasticity (++) (zones 2,3,4,5, and 6). Zones 7 and 8 are located at maximum biomass requirements in the abundance-growth space and exhibit fixed flux values (+/0).

Characterizing *P_F,μ_* for each point of the grid is time-consuming and complex to interpret. Thus, given (*F, μ*), we compute the range [*min_F,μ_*(*r*), *max_F,μ_*(*r*)] of the flux *v_r_* fora reaction *r*, where limits of this interval are the minimal and maximal values of *v_r_* in *P_F,μ_*. This range serves as an indicator of metabolic plasticity. More precisely, given an abundance-growth point (*F, μ*), a reaction r has *flux plasticity* if *min_F,μ_*(*r*) ≠ *max_F,μ_*(*r*). In this case, the reaction also has *structural plasticity* if zero is included in this interval, meaning that alternative pathways can replace *r* (30, 31).

Following this rationale, for studying metabolic phenotypes given by a set of reactions of interest *R′* on an abundance-growth point (*F,μ*), we assign a categorical value to *r* ∈ *R′* according to Table 1. Exchange reactions can be selected as *R′* for focusing on the community’s metabolic interactions. To explore plasticity changes across this space, for each point of the grid, we compute a categorical vector for reactions in *R′*, generating a *partition* of the abundance-growth space in zones of identical categorical vectors. Depending on the size of R’, the number of zones can be huge, making it challenging to analyze. To overcome this, we group points in a *cluster-partition* obtained using a clustering algorithm (Methods). Atoy problem in Figure 1 illustrates a partition of the abundance-growth space of a community of two organisms. Shifts in categorical values along this partition are associated with alternative pathways for biomass production.

**Table 1.** Categorical values for a reaction *r* and a point (*F,μ*) of the abundancegrowth space.

| Condition | Category | Flux sign | Flux variability | Flux requirement | Level of plasticity |
| --- | --- | --- | --- | --- | --- |
| $\min_{F, \mu}(r) = \max_{F, \mu}(r) = 0$ | 0 | zero | fixed | off | no plasticity |
| $\min_{F, \mu}(r) = \max_{F, \mu}(r) < 0$ | – | negative | fixed | mandatory | no plasticity |
| $0 < \min_{F, \mu}(r) = \max_{F, \mu}(r)$ | + | positive | fixed | mandatory | no plasticity |
| $\min_{F, \mu}(r) < \max_{F, \mu}(r) < 0$ | -- | negative | variable | mandatory | flux plasticity |
| $0 < \min_{F, \mu}(r) < \max_{F, \mu}(r)$ | ++ | positive | variable | mandatory | flux plasticity |
| $\min_{F, \mu}(r) < \max_{F, \mu}(r) = 0$ | –0 | negative or zero | variable | optional | structural plasticity |
| $0 = \min_{F, \mu}(r) < \max_{F, \mu}(r)$ | 0+ | positive or zero | variable | optional | structural plasticity |
| $\min_{F, \mu}(r) < 0 < \max_{F, \mu}(r)$ | –+ | any | variable | optional | structural plasticity |

### Analysis of the abundance-growth space for a bioleaching community

A simple microbial consortium is modeling metabolic cross-talking between *A. ferrooxidans* Wenelen and *S. thermosulfidooxidans* Cutipay (File S1, File S2, Methods), obtaining a community competing for inorganic energy sources (Fe(II) and thiosulfate). Only Cutipay can consume organic matter made available by the degradation of community biomass from cellular death, which is modeled as a pseudo-reaction transforming a fraction *α* of the biomass produced by the community (Methods).

We considered the case study when the community is in the presence of organic matter (*α* = 0.2) and a medium substrate level composed of energetically equivalent amounts of Fe(II) and thiosulfate. With *R’* the set of 100 exchange reactions, the abundance-growth space is partitioned into 69 zones. For distinguishing the most relevant changes in this space, a cluster-partition is computed using 20 clusters (Figure 2(b)) (Methods and Figure S1 to justify the choice of the number of clusters).

**FIG 2.**
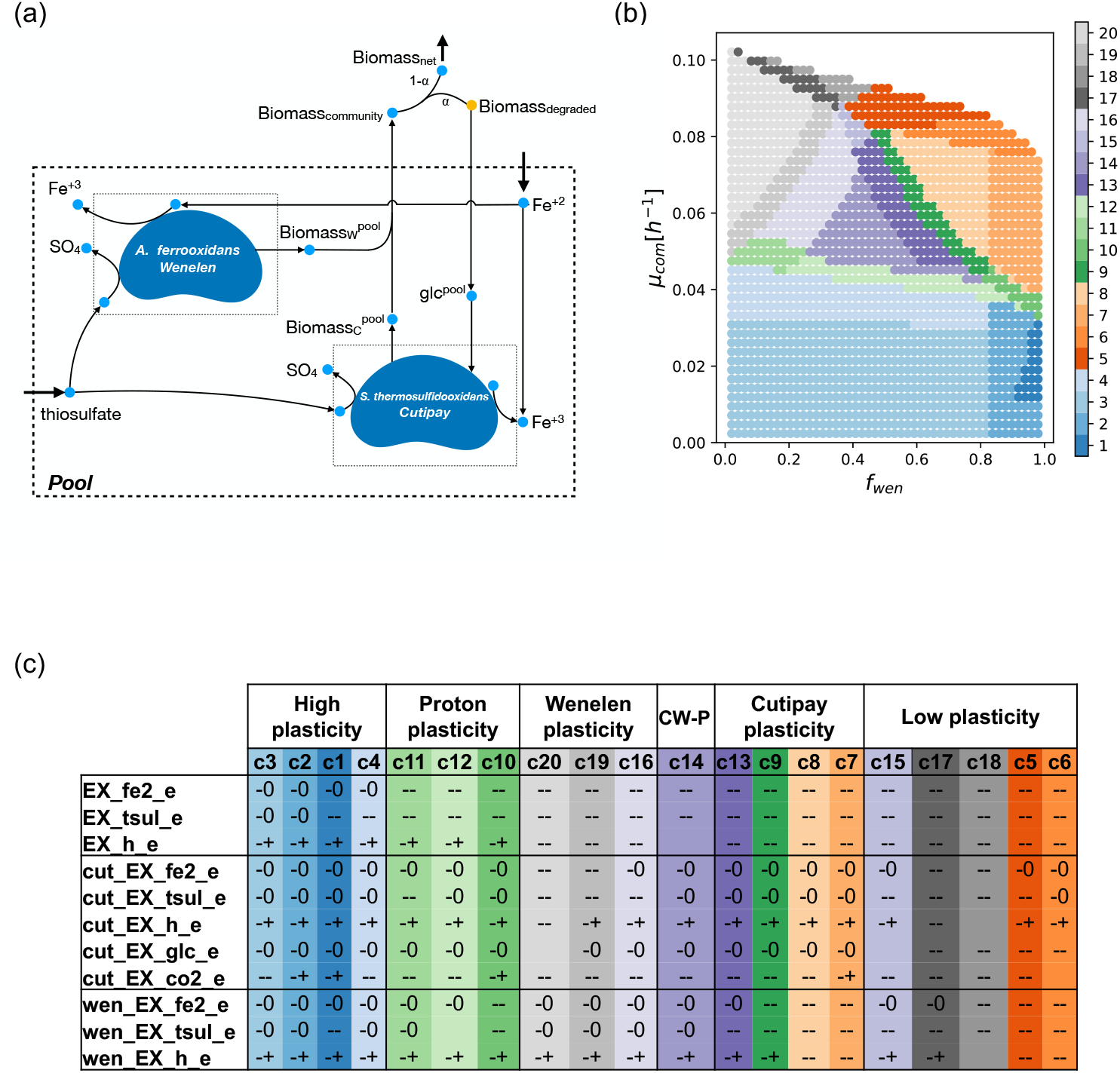
Analysis of the abundance-growth space fora bioleaching community: reference case. (a) A community comprised by *A. ferrooxidans* Wenelen (wen) and *S. thermosulfi-dooxidans* Cutipay (cut) is studied in an environment with Fe(II) and thiosulfate in the presence of organic matter. A pseudo reaction represents organic matter availability as a fraction (*α*) of the total community biomass. (b) Twenty clusters are computed for the 100 exchange reactions of the community model. (c) Key reactions associated with qualitative changes between neighboring clusters define six areas for the behavior of this community. A status is associated with a cluster when over 80% of the points in the cluster exhibit a certain state. If no consensus is reached, an empty cell is presented. Clusters are organized according to their location in the abundance-growth space from lower to high growth rate and from f_*wen*_ 0 to 1. EX: exchange reactions, h: hydrogen (proton), tsul: thiosulfate, co2: carbon dioxide, fe2: Fe(II), glc: glucose, CW-P: Cutipay Wenelen plasticity.

Clusters were analyzed by selecting reactions that display different qualitative states between neighboring clusters, finding that changes in 23 reactions give rise to the 20 clusters observed in this scenario (Figure 2 (b)). Seventy-seven remaining reactions are blocked, coupled with their biomass production requirement, or exhibited structural plasticity on each grid point. The curation of these 23 reactions allowed us to remove reactions coupled in all clusters, such as consumption of Fe(II) and production of Fe(III), pairs of exchange reactions for H_2_O and H, and SO_4_ and H. Additionally, reactions in which qualitative changes are only attributed to a few clusters were removed, notably the exchange of H_2_, in which production is optional only in cluster 1. After this curation, eleven critical reactions associated with iron, sulfur, and carbon metabolism were obtained. Five of them belong to Cutipay, three to Wenelen, and three to the microbial community as a whole (Figure 2 (c)).

Six sections are distinguished (Figure 2 (b) and (c)). A high plasticity section is defined at lower growth rates, where most exchanges are optional (i.e., −0 and -+) and independent of relative abundances. As growth requirements increase, oxidation of both inorganic energy sources by the community becomes mandatory (i.e., the category-), as well as the consumption of sulfur and iron by both bacteria, which indicates a loosing of global structural plasticity. Next, we define a proton plasticity section (clusters 10, 11, and 12), where the community needs to consume Fe(II) and thiosulfate, preserving structural plasticity (-+) for the exchange of protons (H) as a community. Proton exchange indirectly measures the balance between Fe(II) and thiosulfate oxidation since iron oxidation requires protons, while thiosulfate oxidation produces them (32). Thus, in this section, the community can still opt not to affect the pH of its environment.

In the Wenelen plasticity section, all reactions associated with this bacterium exhibit structural plasticity. In this section, Cutipay shifts from no plasticity for Fe(II) oxidation in cluster 20 (where it is abundant) to structural plasticity in cluster 16. In this transition, we observe a switch from status - to -+ for cut_EX_h_e, representing a less active Fe(II) oxidation.

Analogously, the Cutipay plasticity section shows the behavior of all associated reactions, while Wenelen loses its structural plasticity for thiosulfate (cluster 13) and iron (cluster 9) consumption when moving towards *f_wen_* = 1. Cluster 7 shows an increase in carbon metabolism structural plasticity depicted by a shift in the qualitative state of EX_co2_e in Cutipay, where it no longer requires CO_2_ consumption to support its growth.

In between the above two sections stands the Cutipay-Wenelen plasticity section, where high plasticity is observed for both organisms. It is a desirable area where organisms can adapt to shifts in their environment, maintaining high growth rates. Finally, in the low plasticity section, characterized by the highest growth rates, all reactions lose plasticity, and the high energy requirements force Cutipay to consume glucose.

Despite their apparent symmetry, Cutipay can maintain flexibility at higher growth rates than Wenelen. Different efficiencies influence this in metabolizing energy sources and its additional energy and carbon source (glucose). These differences also affect the distribution of maximum growth rates of the community, located in areas in the abundance-growth space where Cutipay is predominant.

Eleven critical exchange reactions were selected from the previous clustering analysis. By performing a partition using these essential reactions, 40 areas are obtained, among which an area where glucose consumption of Cutipay is clearly defined as well as a much specific description of the low plasticity section (Figure 3), thus confirming in a deterministic way previous observations regarding plasticity in the abundance-growth space.

**FIG 3.**
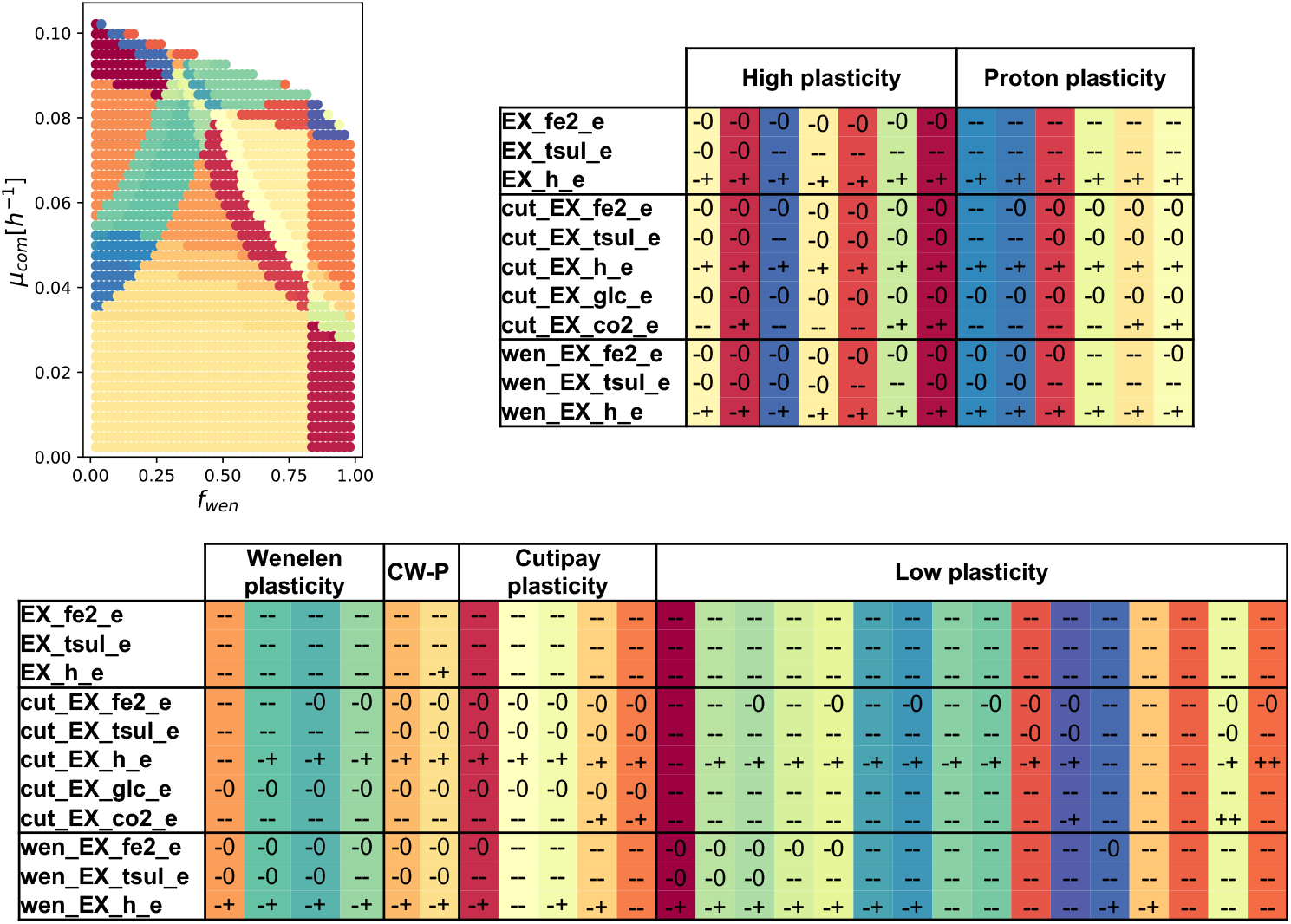
Partition of the abundance-growth space determined by 11 selected key bioleaching reactions describes relevant shifts in the bioleaching community. The partition of the abundance-growth space determined from reactions for oxidation of iron and sulfur and carbon metabolism shows that critical exchange reactions associated with energy resources and carbon metabolism can fully characterize the metabolic shifts occurring in the bioleaching community, yielding 40 areas with defined qualitative status. Reactions with EX prefix denote exchange reactions, cut: *S. thermosulfidooxidans* Cutipay, wen: *A. ferrooxidans* Wenelen, co2: carbon dioxide, fe2: Fe(II), glc: glucose, h: hydrogen, tsul: thiosulfate, CW-P: Cutipay Wenelen plasticity. Partitions are organized according to their location in the abundance-growth space from lower to high growth rate and from f_*wen*_ 0 to 1.

To quantitatively illustrate previous results, explore competition for resources (Figure 4(a), Figure S2) and use of different substrates to support cell growth (Figure 4(b), Figure S3), we computed the feasible solution space of critical reactions associated with consumption of energy sources at specific points of the abundance-growth space.

**FIG 4.**
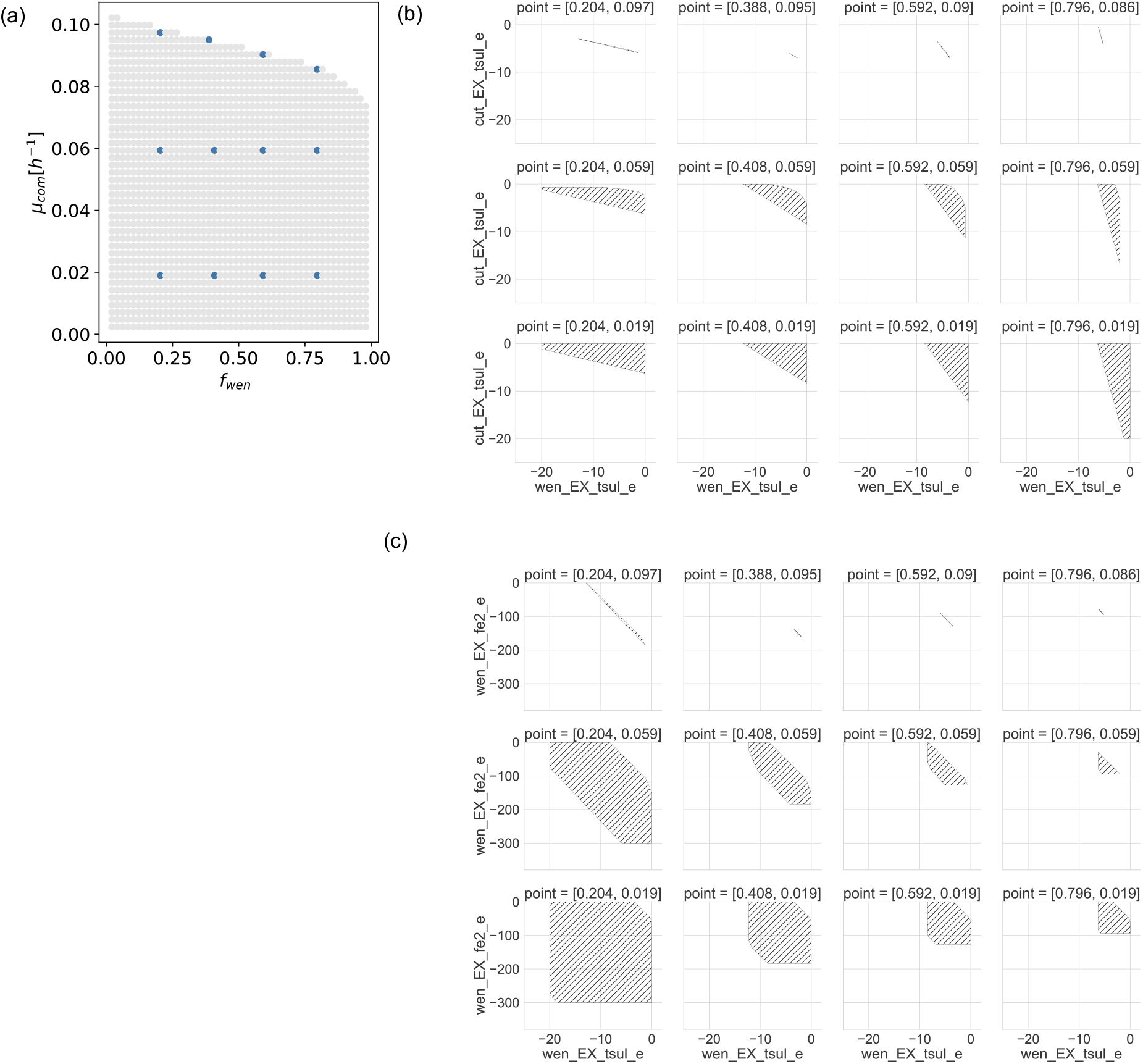
Flux relation analysis. Changes in the feasible fluxes of reactions of interest for *A. ferrooxidans* Wenelen and *S. thermosulfidooxidans* Cutipay computed for specific points in the abundance-growth space (a). Computed reaction fluxes represent competition for thiosulfate for both bacteria (b)and distribution of resources for Wenelen (c). Fluxes are in units of [mmol/gDW_*org*_ h^−1^]

For each analyzed point, the degree of plasticity and relations among reactions are expressed by the shape of the feasible solution space. As expected, there is an overall reduction in plasticity while moving into the growth-oriented area, where exchange fluxes for energy sources are perfectly coupled, exhibiting a functional relation. This result is evident in the point (0.388, 0.095), where for a given consumption of thiosulfate, there is a unique consumption of iron (Figure 4 (b)). Plasticity reduction is also observed when moving into extreme compositions of the community, where being more abundant results in having less specific consumption rates.

### Impact of organic matter availability on a bioleaching community

Organic compounds are detrimental for some autotrophic bioleaching bacteria (33), toxic for chemolithotrophic (34), having inhibitory effects in iron oxidation (35), thus shaping community dynamics in bioleaching communities favoring abundance of heterotrophs or facultative heterotrophs (36).

Different scenarios with increasing amounts of organic matter availability were simulated by moving *α*. For each scenario, partitions of the abundance-growth space were computed for essential bioleaching reactions (Figure 5). By increasing *α*, we observe a direct effect in the enlargement of the space with the maximum growth rate of the community. Additionally, an increase of the higher structural plasticity areas is observed (Figure 5), but in Wenelen, this only happens until the maximum uptake of inorganic sources is reached (*α* = 0.6). Interestingly, in Cutipay, the enlargement of the space is given by the mandatory glucose consumption (i.e., - - state), and it exhibits a shift where CO_2_ consumption is no longer required to support its growth (a grey area in Figure 5 (a)). Moreover, new areas appeared near the maximum community growth section, where this bacterium is forced to break down glucose, hence producing CO_2_ in order to survive (Figure 5). These initial results show that organic matter is beneficial for achieving higher growth rates and that Cutipay does not present a preference for glucose catabolism over oxidation of inorganic sources.

**FIG 5.**
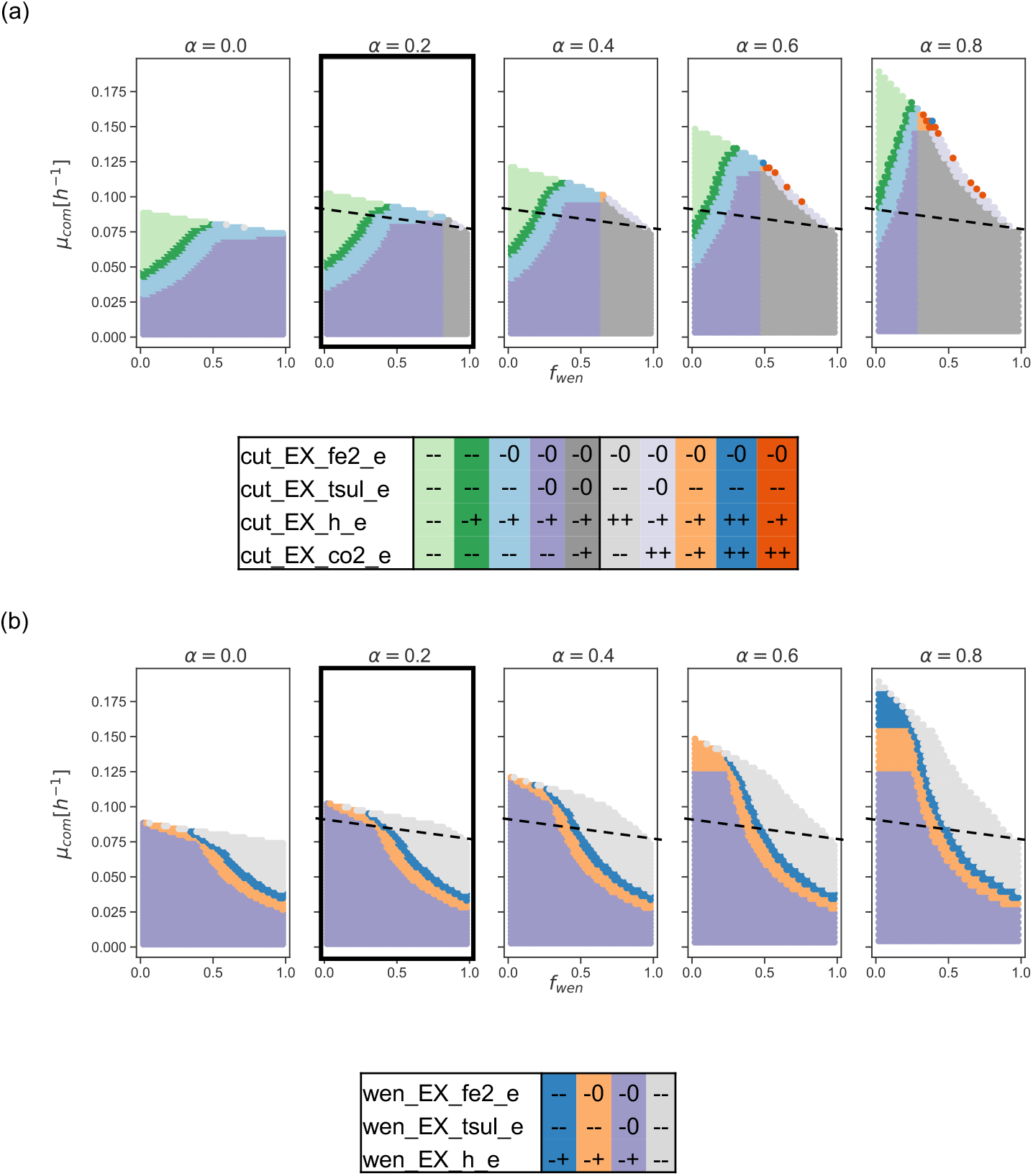
Effects of organic matter availability. A partition of the abundance-growth space is computed for key reactions for *S. thermosulfidooxidans* Cutipay (a) and *A. ferrooxidans* Wenelen (b) while changing the organic matter availability (*α*) on each column. *α* represents the ratio of the biomass generated by the community that could be degraded to serve as an additional carbon source. The reference case is highlighted with thicker edges; dashed lines show the maximum community growth rate for *α* = 0, over which Cutipay is forced to consume glucose. EX_fe2, EX_tsul: exchange reactions for Fe(II) and thiosulfate, respectively. cut_EX_co2, cut_EX_h: exchange reactions for Cutipay for CO_2_ and H respectively; wen_EX_fe2, wen_EX_tsul, wen_EX_h: exchange reactions for Wenelen for Fe(II), thiosulfate and H. Partition tables are ordered according to the location of different zones in the abundance-growth space from lower to higher growth rates, from f_*wen*_ 0 to 1, and their occurrence in different scenarios from left to right.

This result, in an unconstrained context, initially contradicts literature regarding the behavior of bioleaching communities in presence of organic matter, which are characterized by an increased abundance of heterotrophs and facultative heterotrophs which favor consumption of carbon sources for energy production (36), and will be explored by imposing additional objectives.

### Impact of substrate availability and composition on the bioleaching community

The effect of substrate availability on the abundance-growth space is studied by changing two external conditions: amounts of Fe(II) and thiosulfate availability and the ratio between these energy sources. For studying the effect of the available energysource, we consider three levels (low, medium, high) of both Fe(II) and thiosulfate, with *α* = 0.2 (Methods). When changing scenarios, partitions of the abundance-growth space built from critical reactions exhibit an increment of the feasible space. Consequently, there is also an increase in the high plasticity areas and maximum community growth rates (Figure 6), which are similar to the findings obtained when performing this analysis with *α* = 0.6 (Figure S4).

**FIG 6.**
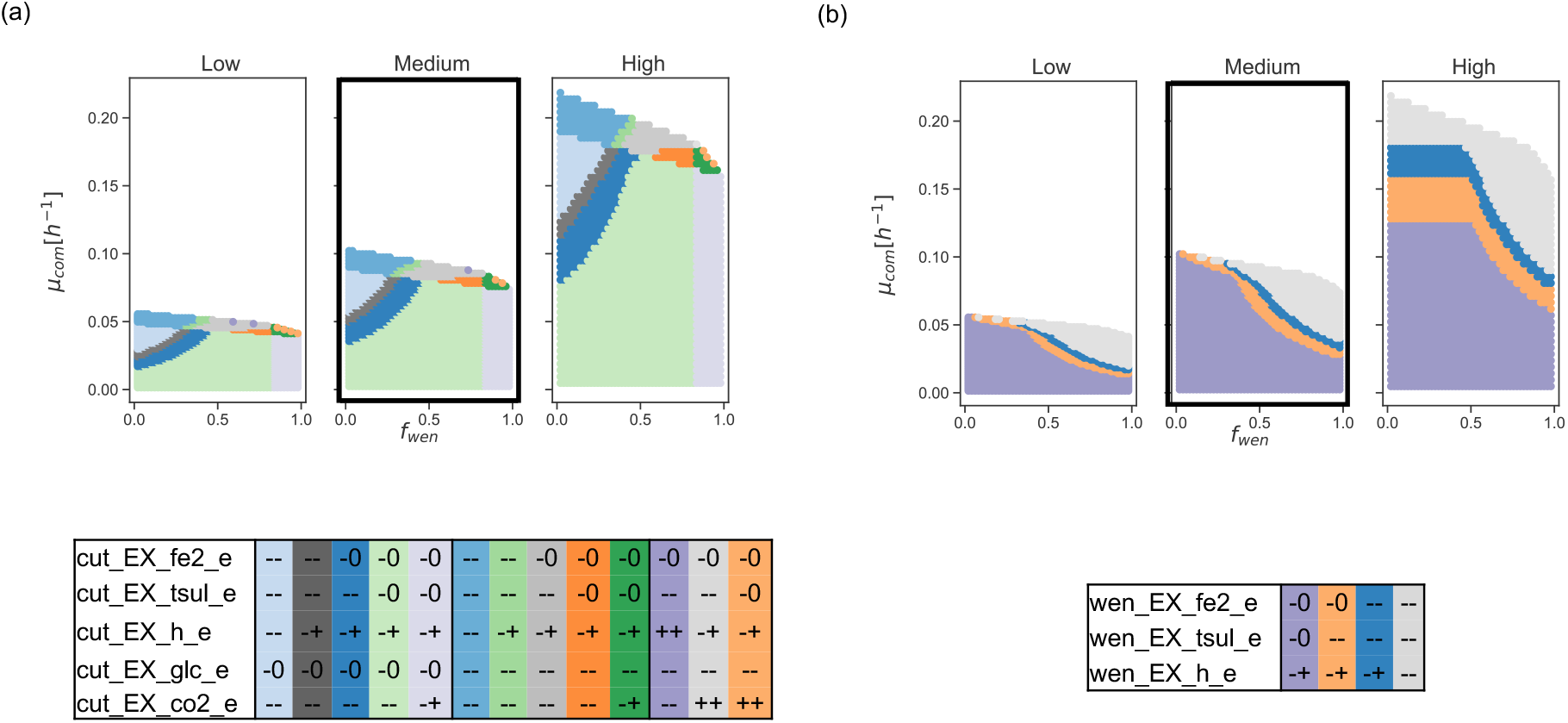
Effect of substrate availability. A partition of the abundance-growth space defined by qualitative states for key reactions of *S. thermosulfidooxidans* Cutipay (a) and *A. ferrooxidans* Wenelen (b) is presented in scenarios with changing availability of Fe(II) and thiosulfate. Three levels of availability are studied, where the reference scenario is the average case. The reference case is highlighted with thicker edges. Partition tables are ordered according to the location of different zones in the abundance-growth space from lower to higher growth rates, from f_*wen*_ 0 to 1, and from their occurrence in different scenarios from left to right.

Different stages of the bioleaching process are characterized by changes in chemical and physical properties, which shape the distribution of resources, metabolic states, and community compositions (36). In particular, uniform distribution of substrates is not always guaranteed, and nutrient bioavailability could pose a bottleneck for bioleaching efficiency.

Consistently, in our model, changing the relative availability of Fe(II) and thiosulfate has drastic effects on the behavior of the community (Figure 7). Iron-predominant scenarios present new metabolic states where iron availability overpasses the effect of preference for thiosulfate as an energy source, which has been observed up to this point in these analyses. It is worth noticing that sulfur compounds have higher energy yields than Fe(II) since they have more electrons available (37). Although, *A. ferrooxidans* is believed to exhibit a preference for iron (38), evidence shows the simultaneous expression of genes forconsumption of iron and sulfur (39), being solubilization of sulfur compounds key to consuming both energy sources (40). This result is consistent with our analysis, where a preference for Fe(II) oxidation is only displayed when its availability surpasses thiosulfate.

**FIG 7.**
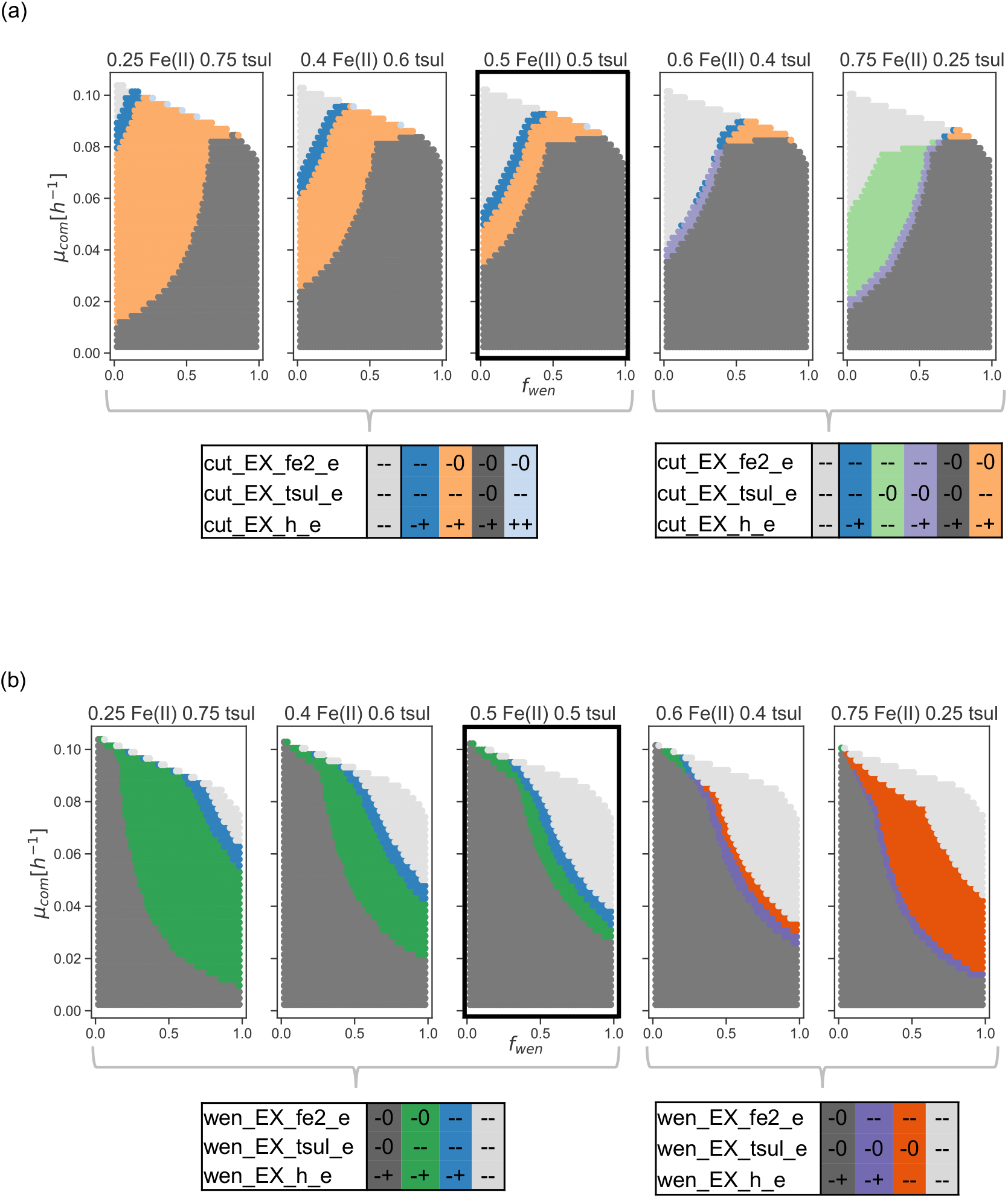
Effect of relative substrate availability. A partition of the abundance-growth space defined by qualitative states for key reactions of *S. thermosulfidooxidans* Cutipay (a) and *A. ferrooxidans* Wenelen (b) is presented in scenarios with medium substrate availability but changing ratios between both energy sources. The reference case is highlighted with thicker edges. Two partition tables for each bacteria are displayed for the qualitative states in different scenarios. Partition tables are ordered according to the location of different zones in the abundance-growth space from lower to higher growth rates, from f_*wen*_ 0 to 1, and from their occurrence in different scenarios from left to right.

### Impact of considering alternative objectives

For testing a more realistic representation of a bioleaching community, two additional metabolic objectives were selected and simulated separately: a parsimonious approach, where the sum of the overall fluxes is minimized, and the maximization of extracellular polymeric substances (EPS) to represent bioleaching bacteria attached to a mineral.

A parsimonious FBA, where minimization of the sum of fluxes is performed as a proxy for energetic efficiency, has surged as a realistic alternative to retrieve flux distributions (41, 42). This analysis allows a 10%, 20%, and 50% deviation from the optimal solution to maintain a certain degree of plasticity. The results (Figure 8) show a significant change with the one observed in Figure 3.

**FIG 8.**
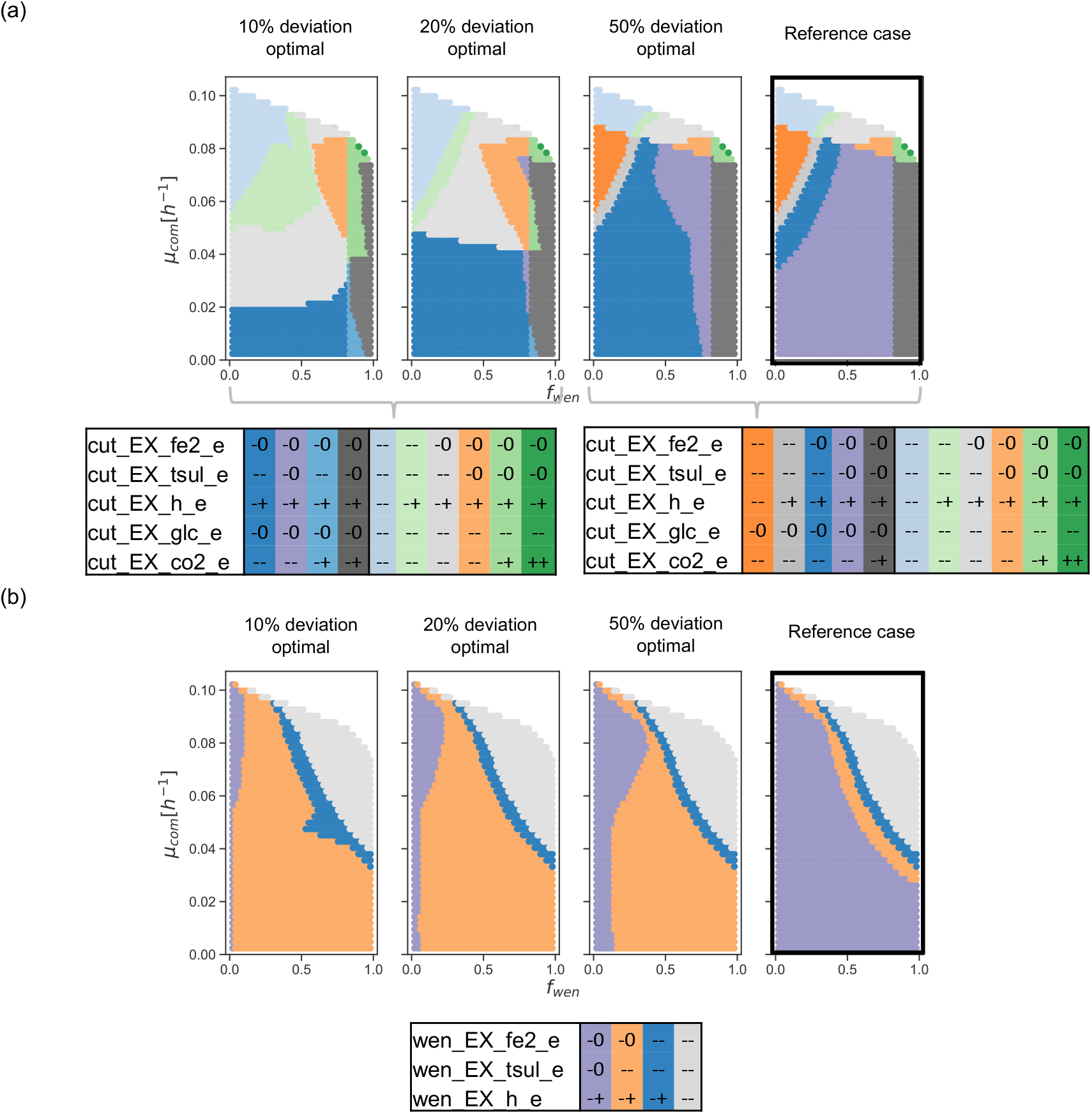
Effect on qualitative states by alternative objective functions - energetic efficiency. A partition of the abundance-growth space for key reactions of *S. thermosul-fidooxidans* Cutipay (a) and *A. ferrooxidans* Wenelen (b) is depicted in a scenario where 10%, 20%, and 50% of deviation from the optimal solution for energy efficiency is allowed. Two partition tables are displayed for Cutipay for the qualitative states presented in different scenarios. Partition tables are ordered according to the location of different zones in the abundance-growth space from lower to higher growth rates, from f_*wen*_ 0 to 1, and from their occurrence in different scenarios from left to right.

Regarding energy source preferences, it is remarkable how thiosulfate becomes mandatory at significantly lower growth rates as we are close to the optimal energy efficiency value. However, the degrees of plasticity in Fe(II) consumption remain unaltered concerning the reference case in both bacteria. This observation is valid in all areas of the space, except for Cutipay in *f_wen_* ≥ 0.8, where this bacteria retains structural plasticity for both inorganic sources. Interestingly, glucose consumption by this bacteria is required at lower growth rates, which is in agreement with the literature, where the presence of organic matter is beneficial for heterotrophs that use this energy source for biomass production (36, 43, 44).

Metabolic states in bioleaching bacteria depend highly on their growth conditions, particularly if they are attached to the mineral or growing in a bioleaching solution (45). This adherence to the mineral is achieved by synthesizing extracellular polymeric substances (EPS) that provide an extracellular matrix for attachment and concentration of ions, creating a beneficial environment for bioleaching efficiency (45, 46, 47).

Synthesis of EPS by the community is represented as an additional chemical reaction that either bacteria can perform. A description of the abundance-growth space is obtained considering the production of at least 20%, 50%, and 90% of the maximum EPS produced at each point of the space.

Concerning energy source preferences, Cutipay changes its substrate preference, mainly relying on glucose and thiosulfate when approaching maximum production of EPS (90%) ((Figure 9) (a)). However, for Wenelen, there are no significant changes in plasticity except near optimal EPS production, and where its abundance exceeds 80%. It is where this bacteria require all energy sources.

**FIG 9.**
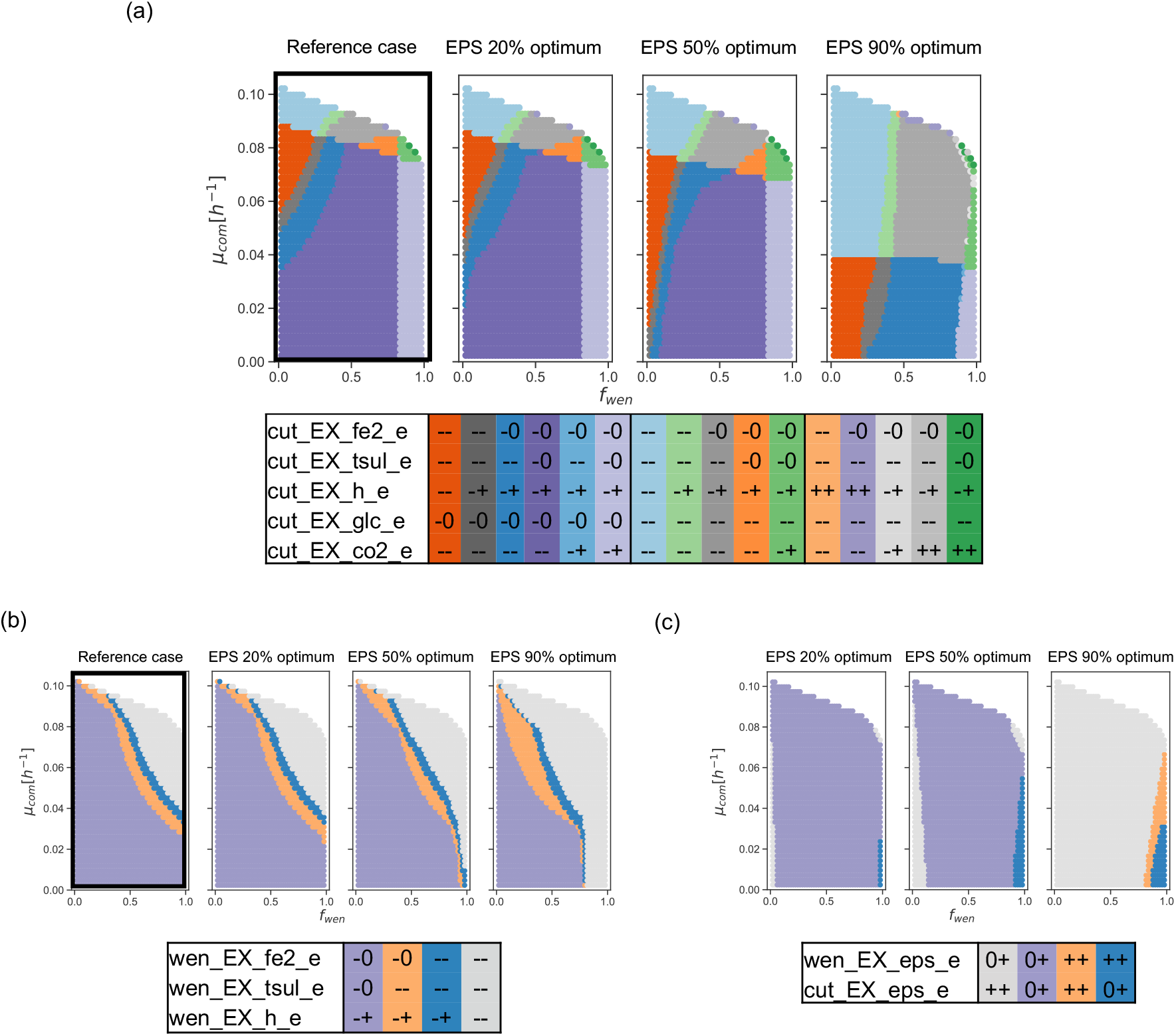
Effects on qualitative changes by alternative objective functions - EPS production. (a) partition of the abundance-growth space for *S. thermosulfidooxidans* Cutipay key exchange reactions, (b) partition of the abundance-growth space for *A. ferrooxidans* Wenelen and (c) partition for EPS production by both bacteria in the bioleaching community. The reference case is highlighted with thicker edges. Partition tables are ordered according to the location of different zones in the abundance-growth space from lower to highergrowth rates, from f_*wen*_ 0 to 1, and from their occurrence in different scenarios from left to right.

Although both members of the community are modeled to synthesize EPS potentially, only Cutipay is forced when approaching the maximal production (Figure 9 (c)). It contradicts experimental data showing that this bacteria is a weak biofilm performer (48, 49). However, it has been reported that *S. thermosulfidooxidans* improves its EPS production capacity after replenishing growth media, given by the nutritional boost of media replacement (49), a metabolic state represented in these simulations. Additionally, synthesizing compounds comprising Cutipay biomass requires less energy than Wenelen, which puts Cutipay at an advantage in synthesizing this extracellular polymer.

## DISCUSSION

Predicting the metabolic phenotype of a microbial community in a given environment is a challenging problem. Constraint-based modeling for single organisms has historically based most of their flux predictions on the maximization of biomass production (50). This assumption yields promising results for organisms well characterized in laboratory settings where they have been isolated for being efficient in the growth and synthesis of a product of interest due to their use in biotechnological applications (51, 52). In contrast, organisms in nature are exposed to constant shifts in their environment, which implies, in particular, the production of secondary metabolites to provide favorable conditions for survival, deviating resources to the detriment of a growth-driven metabolism (5). The nature of their gives increased complexity of inter-species interactions (53, 54), where cross-feeding between organisms gains relevance and the composition of a community, given by the relative abundances or the organisms that comprise it. Thus, establishing metabolic interactions in nature is a challenging task, and as a consequence, predicting flux distributions, or metabolic phenotypes, requires new strategies.

Due to the increasing interest in studying communities, several approaches have been developed to characterize interactions between organisms (12, 14) and model their behavior in different case studies relevant to the biotechnological industry (13, 15). From a more ecological perspective, an innovative notion of the metabolic niche was proposed to characterize all possible environments in which an organism could live in sub-optimal conditions (i.e., no imposition of maximizing an objective function) (28). This adaptation of the ecological niche concept uses the feasible flux space to study how organisms interact with their environment, represented by few exchange reactions, and was tested by comparing metabolic niches for over 500 microorganisms to provide a new niche characterization that depicts the relationship between genotype and phenotype (28). Even though this method explores all feasible metabolic fluxes for a single organism without the imposition of an objective function, it does not fully characterize emergent properties arising from a specific community composition.

In this work, we introduce the abundance-growth space defined by the composition of a community and its growth rate as an approach to characterize the space of feasible metabolic fluxes in a community. More precisely, we partition the abundance-growth space based on the flux variability of a relevant set of reactions compatible with each one of the points of this space. By performing this analysis, we can find metabolic phenotypes that characterize different areas of this space and address the question of how plasticity, represented as a measure of the range of feasible fluxes in a point for a specific reaction, changes across suboptimal growth rates relevant for the study of natural communities. Additionally, this method explains how relations between reactions change in different suboptimal growth states along the abundance-growth space.

Environmental communities are subjected to constant shifts that are detrimental to their survival. Adaptation to such changes include gene mutations (54), gene transfer, and even gene loss in collaborative communities (53), as well as the development of regulatory processes that result in efficient distribution of resources such as flux distributions that result on increased metabolic plasticity (55). This raises the long-standing question of whether the community *opts* to live at suboptimal growth rates to increase the capacity to use alternative metabolic pathways (42). By studying a bioleaching community, this method allows for unveiling how plasticity changes in the abundance-growth space, confirming that maximum plasticity is attained at lower growth rates. Moreover, the method confirms that by increasing the growth rate of the community, the relative abundance of its members is relevant, and the coupling of reactions becomes frequent, rigidifying their metabolism.

The clustering of points in the abundance-growth space, according to their plasticity, allows us to pinpoint critical reactions involved in significant changes in interactions of the community with the environment and establishes in a precise manner where critical changes in plasticity occur. Notably, fora bioleaching community, few reactions are found to explain most of their changes in plasticity, and the composition of available energy sources is critical for community plasticity. Additionally, by considering either energetic efficiency or the production of extracellular polymeric substances, the method retrieves a preference for carbon sources, which coincides with literature observations, and a detriment in plasticity for inorganic energy sources.

However, the characterization of reactions based on their plasticity does not allow for the recovery of the degree of coupling between reactions. This calls for performing a flux relation analysis, illustrating how the coupling between reactions changes along the abundance-growth space. In the example developed in this paper, this was illustrated with the consumption of energy sources, showing that despite their exhibited plasticity, there are strong couplings when reaching high growth rates. In this work, we proposed a novel framework for studying the catalog of different metabolic phenotypes that a community could display from a by-product-driven metabolism towards a growth-driven metabolism which is usually represented using other methods for studying microbial communities (15, 13, 16, 17). This adaptation of the metabolic niche could serve as a framework to study environmental perturbations and the effect of gene perturbations in communities, hence being one step further to unveiling drivers for interactions in ecological communities.

## MATERIALS AND METHODS

### Modeling microbial community metabolism

An expansion of constraint-based modeling for microbial communities was proposed by (13). In summary, *N* organisms belonging to a community are considered. For each organism *i* ∈ {1,…,*N*}, a single model comprised a set of *M_i_* metabolites differentiated by organism and compartment and a set *R_i_* of all reactions of organism *i*. These reactions include all internal reactions, transport between compartments, and exchange of metabolites with their media. A biomass reaction *r_biom_i_* is also included in R, whose stoichiometric values on their substrates correspond to the amounts (in *mmol*) of metabolites present in one gram of dry weight of biomass of the organism *i* (*gDW_i_*).

If *v_r_* denotes the flux over reaction *r* expressed in [*mmol/gDW_i_/h*], then the following constraints exist for organism *i*:

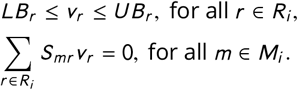

The constants *LB_r_* and *UB_r_* are the lower and upper bounds for the flux over a reaction *r* and are used to control the amount and direction of fluxes in the system.

Given *N* single-organism models, a community is represented as a single compartmentalized system as previously established in (16, 56). In this model, each metabolic network is incorporated as a meta-compartment with an additional *pool* compartment that represents the shared media where metabolites can be exchanged among community members and the exterior.

A set of *pool metabolites M_pool_* is created containing metabolites present in the extracellular compartment of at least one organism. Consequently, any exchange reaction of organism *i* is considered a transport reaction between the extracellular compartment of *i* and the pool. Finally, a new set *R^EX^* of exchange reactions is defined to control external conditions for all metabolites in the pool.

In this community model, each member *i* makes up a fraction *f_i_* of the total amount of community biomass (with 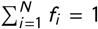). These fractions are relevant to compare the feasible fluxes on reactions of different organisms accurately.

Indeed, in this community model, fluxes are expressed in a single unit [*mmol/gDW_com_* h^−1^], where *gDW_com_* is a gram of dry weight of the *community biomass*. Since bounds on reaction fluxes of organism *i* were originally expressed in [*mmol/gDW_com_* h^-1^] in the single-organism model, they must be recomputed in the community model, which is done by weighting by *f_i_* each bound previously defined on a flux reaction of organism *i* (15).

Given this unit conversion, the flux *v_biom_i_* of the biomass reaction of organism *i* is not expressing the growth *μ_i_* of organism *i* but the grams of organism *i* produced by each gram of the *community* per hour. Hence, if we consider a state of balanced growth, where the fraction *f_i_* of each organism *i* is maintained over time, then we should impose that all organisms are growing at the same rate: *μ*_1_ =…= *μ_N_* = *μ_com_*, which is equivalent to imposing: *v_biom_i_* = *f_i_ μ_com_* for all *i* ∈ {1,…,*N*}(15).

Considering all these observations, the set of constraints on the fluxes *v_r_* of the community are the following:

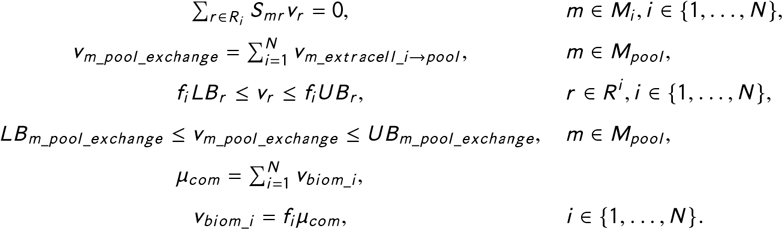

If the fractions *f_i_* are fixed (as in cFBA (16)) or if the community growth rate *μ_com_* is fixed (as in the case of SteadyCom (15) and RedCom (13)), then all constraints are linear.

Eventually, any other interaction between the organisms or additional information that can be expressed as a linear constraint on the fluxes can be easily added to the model. For instance, the model can directly include fixing values to reaction fluxes or imposing coupling between reactions. In the case of the bioleaching community presented in this work, a pseudo-reaction models the capacity of one organism to use degraded organic matter as an additional energy source.

### Analysis of the abundance-growth space

The definition and analysis of the abundance-growth space are performed as follows.

#### Definition of abundance-growth space

Considering *N* organisms in the community, an *abundance-growth point* is given by 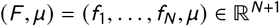, where *f*_1_,…,*f_N_* are the relative abundances of the organisms (with 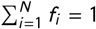) and *μ* is the growth rate of the community (15).

An abundance-growth point (*F,μ*) is *feasible* if given *F* and *μ*, at least one flux distribution satisfies all constraints of the community model.

#### Building a grid of feasible abundance-growth points

A set 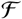 is built, corresponding to a discretization of all possible abundance-growth points. Specifically, for a given resolution parameter ℓ, we define a discretization of the possible relative abundances values by consideringthe set 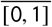 corresponding to ℓ equidistant values in the interval [0,1]. Thus, the set 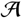 of relative abundance points (*f*_1_,…,*f_N_*) that we consider in the analysis are:

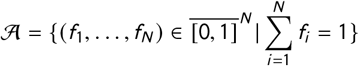

A second discretization is done for the community growth rate *μ* by defining the set 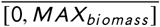 corresponding to ℓequidistant values in the interval [0, *MAX_biomass_*], where *MAX_biomass_* is the maximum biomass computed in the model for all abundance points in 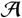. The set of abundance-growth points considered in the analysis is defined by:

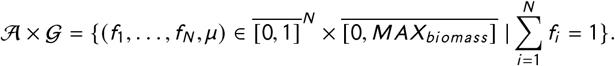

Finally, the set of points that define the grid 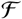 are those that are feasible. That is, the points (*f*_1_,…,*f_N_*,*μ*) in 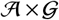 such that the community can reach a growth *μ* for the given relative abundances (*f*_1_,…,*f_N_*).

Note that, since 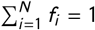 then an abundance of one organism can be omitted in an abundance-growth point, which is especially useful in the case when *N* = 2, since this implies that abundance-growth points (*f*_1_, *μ*) can be depicted in a two-dimensional plot.

#### Computing qualitative vectors for the abundance-growth points

Each point (*F, μ*) of the abundance-growth space is a representation of all the flux distributions in its associated polytope *P_F,μ_*. Given a point 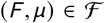 in the abundance-growth space, the range of feasible fluxes of a reaction *r* is exactly the interval [*min_F,μ_*(*r*), *max_F,μ_*(*r*)], where *min*(*r*) = *min*{*v_r_* | for all *v* ∈ *P_F,μ_*}, *max*(*r*) = *max*{*v_r_* | for all *v* ∈ *P_F,μ_*}. Both amounts are easily determined in the community model by running Flux Variability Analysis (FVA) (27) fixing *F* and *μ* in the constraint-based model. Thus, the interval [*min_F,μ_*(*r*), *max_F,μ_*(*r*)] is computed for all reactions *r* in each point (*F,μ*) of the grid 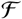. Computation of these amounts can be done in parallel or by efficiently implementing consecutive LP formulations with the same feasible space (57). The range [*min_F,μ_*(*r*), *max_F,μ_*(*r*)] is used to give a categorical value to each reaction *r* and abundance-growth point (*F,μ*) (Table 1). These qualitative values are encapsulated in the vector 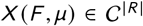 of dimension equal to the number of analyzed reactions in the community model, where 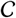 is the set of categories. In the applications of this method, a subset *R′* ⊂ *R* of *selected reactions* will be considered, and thus we will focus our analysis on the restriction of *X*(*F,μ*) to the coordinates associated with *R′*, say 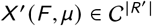.

#### Partition and cluster-partition of the abundance-growth space

As stated previously, the vectors 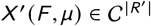 are qualitative descriptions of the set of reactions on each point (*F,μ*) of the grid 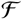. We define two ways of partitioning this grid. First, using the categorical vectors *X′*(*F,μ*), a partition of the grid can be defined by stating that two feasible abundance-growth points in the grid (*F*_1_,*μ*_1_) and (*F*_2_,*μ*_2_) are equivalent if *X′*(*F*_1_,*μ*_1_) = *X′* (*F*_2_,*μ*_2_). Thus, each set of the partition can be considered a zone where selected reactions in the model have identical categorical descriptions. It is called the *partition* of the abundance-growth space defined by the selected reactions in *R′*.

Depending on the size of the set of selected reactions *R′*, the number of zones in this partition can be huge. To address this issue, we propose to use a coarser partition based on the degree of similarity of the categorical vectors. More precisely, we perform a classification of the vectors *X′*(*F, μ*) for each 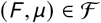 and *r* ∈ *R′* using a hierarchical clustering algorithm with a Jaccard distance, where the number of clusters (*k*) is defined *a priori*. In other words, we can identify zones of the grid where reactions have a similar categorical behavior according to the clustering method used. We call this partition the *cluster-partition* of the abundance-growth space (defined by the selected reactions in *R*’). Alternatively, this method can be implemented using other clustering methods.

Additionally, we want to summarize the qualitative state for each reaction on a given zone defined by a cluster partition. To achieve this, for each reaction, given a cluster, we assign a qualitative value to each reaction if such a qualitative state is present in over 80% of the points of that cluster.

#### Determination of the number of clusters for analysis

Determination of the number of clusters to be used is performed as follows: for a given number *k* of clusters, each categorical vector *X′*(*F,μ*) is compared to its assigned cluster-partition descriptor. For each reaction, a score value between 0 and 1 represents the fraction of points consistent with the cluster-partition descriptor. This analysis is performed with different values of *k* clusters to retrieve the minimum number of clusters where all analyzed reactions are represented correctly in at least 90% of points of the grid.

#### Flux relation analysis

Given an abundance-growth point, the space of feasible fluxes for two given reactions *r*_1_ and *r*_2_ is computed by defining a homogeneous grid of 50 values between the minimum and maximum value of the flux of *r*_1_ computed by FVA. Then, for each flux value of *r*_1_ in this grid, an FVA is performed for reaction *r*_2_ to compute the minimum and maximum flux values of *r*_2_. The interval defined by these values is the feasible fluxes of *r*_2_ for the given flux value of *r*_1_. Normalization by *f_i_* is performed to retrieve flux values in units of [*mmol/gDW_i_/h*] for each analyzed reaction.

### Analysis of bioleaching community

#### Reconstruction of genome-scale models for bioleaching organisms

Two genome-scale models were reconstructed for *Acidithiobacil-lusferrooxidans* Wenelen (iML510) and *Sulfobacillus thermosulfidooxidans* Cutipay (ISM517). Genome sequences for *A. ferrooxidans* Wenelen and *S. thermosulfidooxidans* Cutipay were retrieved from (58, 59, 60). The AuReMe workflow (61, 62) was used for the independent reconstruction of each metabolic network. For *S. thermosulfidooxidans* Cutipay, a model previously reconstructed in (61) for this bacteria, was used as a starting point for manual curation.

For *A. ferrooxidans*, a draft reconstruction was first obtained using reference metabolic models from other species and gene orthology (63). For *A. ferrooxidans* Wenelen, the model iMC507 for *A. ferrooxidans* ATCC23270 (64) was used as a reference (65).

Drafts for both *A. ferrooxidans* Wenelen and *S. thermosulfidooxidans* Cutipay were manually curated. Particular attention was given to iron, and sulfur oxidation metabolism (66, 67). Pathways from these subsystems were completed after including reactions from the literature (File S1) (66, 67, 68, 69). Since no biomass composition information was available for either of the bacteria studied, adaptations of biomass functions from models iMC507 and iHN637 were used for Wenelen and Cutipay, respectively. Both models were checked using Memote to ensure their quality before publication (70, 71).

Final metabolic models for *A. ferrooxidans* Wenelen (iML510) and *S. thermosulfidooxidans* Cutipay (iSM517) are available at the following address: https://github.com/mathomics/ecosystem and in File S2. The obtained models display the associations between 495 genes for *S. thermosulfidooxidans* and a metabolic network comprising 985 metabolites and 1056 reactions. On the other hand, the genomescale model for *A. ferrooxidans* includes 506 genes associated with 612 reactions and 579 metabolites.

Single genome-scale models for *A. ferrooxidans* Wenelen and *S. thermosulfidooxidans* Cutipay were adjusted to exhibit realistic growth rates in simulations with FBA under the presence of iron (Fe(II)) or sulfur (thiosulfate) as energy sources. For this purpose, lower bounds associated with exchanging reactions for Fe(II) and thiosulfate were modified to retrieve growth rates reported in the literature, particularly a maximum growth rate of 0.15 [h^−1^] (64, 43, 72).

#### Metabolic model of the bioleaching community

The individual models previously reconstructed were merged to represent a bioleaching community model in which there is competition for carbon dioxide consumption as well as two external resources: iron (Fe(II)) and sulfur (thiosulfate). Single entry fluxes for these compounds are supplied in a pool compartment to which both bacteria have access and represent their growth environment. The total availability of these resources was modeled by setting the lower bound of the exchange reactions of the community (EX_fe2_e, EX_tsul_e) depen-ding on different analyzed scenarios described in the next section. Other nutrients required for growth were modeled as equally available for both bacteria.

The capacity of Cutipay to use the degraded organic matter of the community as an additional energy source is modeled as an incorporation of the following pseudo reactions representing the degradation of community biomass and its transformation into carbon equivalent units of glucose:

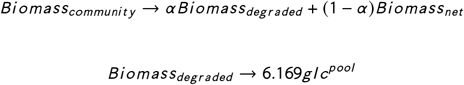

where parameter *α* represents that the death rate of the organisms is equivalent to a fraction *α* of the community growth rate.

The set of 1,717 reactions of this community model comprises 1,668 reactions from the original single models and 49 reactions that appear in the construction of the community model. The set of 1,611 metabolites of the community model comprises 1,564 metabolites from single models and 47 metabolites from the community construction.

#### Definition of parameters and scenarios for simulations of bioleaching communities

Different external and internal conditions are explored to determine how these scenarios change the distribution of qualitative states in the abundance-growth space. External conditions are represented by available resources, both organic and inorganic, for the community. Meanwhile, internal conditions correspond to alternative requirements for the obtained flux distributions, mainly: producing a certain level of EPS or being energetically efficient.

Modeling the availability of organic matter is achieved by setting the parameter *α*. The availability of inorganic sources is modeled, defining the available fluxes of Fe(II) and thiosulfate in the community. It is achieved by adjusting the lower bounds for the community exchange reactions of both sources.

Thus, we define the values Max_Fe(II)_ = 300 [mmol/g DW_*com*_ h^−1^] and Max_tsul_ = 20 [mmol/gDW_*com*_ h^−1^] as reference values, which correspond to the uptakes of Fe(II) and thiosulfate that produce the maximum level of biomass on both organisms according to their single models. Hence, given a parameter *β* ∈ [0, 1] of *energy level* we consider the amounts *β*Max_Fe(II)_ and *β*Max_tsul_ as energetically equivalent.

Any convex combination of these amounts generates an equivalent amount of inorganic sources. Formally, given a energy level *β*, a relative combination of *λ* Fe(II) and 1 − *λ* thiosulfate means that there is a flux of *λβ*Max_Fe(II)_ of Fe(II) and a flux of (1 − *λ*)*β*Max_tsul_ of thiosulfate available to the community. With these definitions, the following scenarios are defined:

1. A *reference case* is defined as having a low organic availability (*α* = 0.2), medium substrate availability (*β* = 0.5) and a relative combination of 0.5 Fe(II) and 0.5 thiosulfate (*λ* = 0.5).
2. Analysis of the effects of organic matter is performed by changing *α* in the reference case scenario (*β* = 0.5, *λ* = 0.5). The analyzed values of *α* are 0, 0.2, 0.4, 0.6, and 0.8.
3. Analysis of the effect of substrate availability in the reference case (*α* = 0.2, *λ* = 0.5) is performed by changing the parameter *β*. The analyzed values of *β* are 1 (high), 0.5 (medium), and 0.3 (low).
4. Analysis of the effect of substrate composition is performed by variations in the value of *λ* in the reference case (*α* = 0.2, *β* = 0.5). The analyzed values of *λ* are 0.25, 0.4, 0.5, 0.6, and 0.75.
5. Analysis of alternative objective functions is performed by analyzing the reference case (*α* = 0.2, *β* = 0.5, *λ* = 0.5) when additional functions are required. Specifically, energetic efficiency is represented as an additional constraint that states that the sum of fluxes of all reactions is less than a factor of the minimum sum of fluxes. The analyzed values for this factor are 1.1,1.2, and 1.5.
6. Analysis of a requirement for EPS production is represented considering the reference case (*α* = 0.2, *β* = 0.5, *λ* = 0.5) with an additional requirement for EPS production. This requirement is represented as an imposition of synthesis of this metabolite as a fraction of the maximum EPS production achievable on each grid point. The analyzed values for this fraction are 0.9, 0.5, and 0.2.

## Availability of data and materials

This method was implemented in Python using COBRApy for constructing community models, FBA, and FVA simulations (73), and it is available at https://github.com/mathomics/ecosystem.

## SUPPLEMENTAL MATERIAL

**FIG S1.**
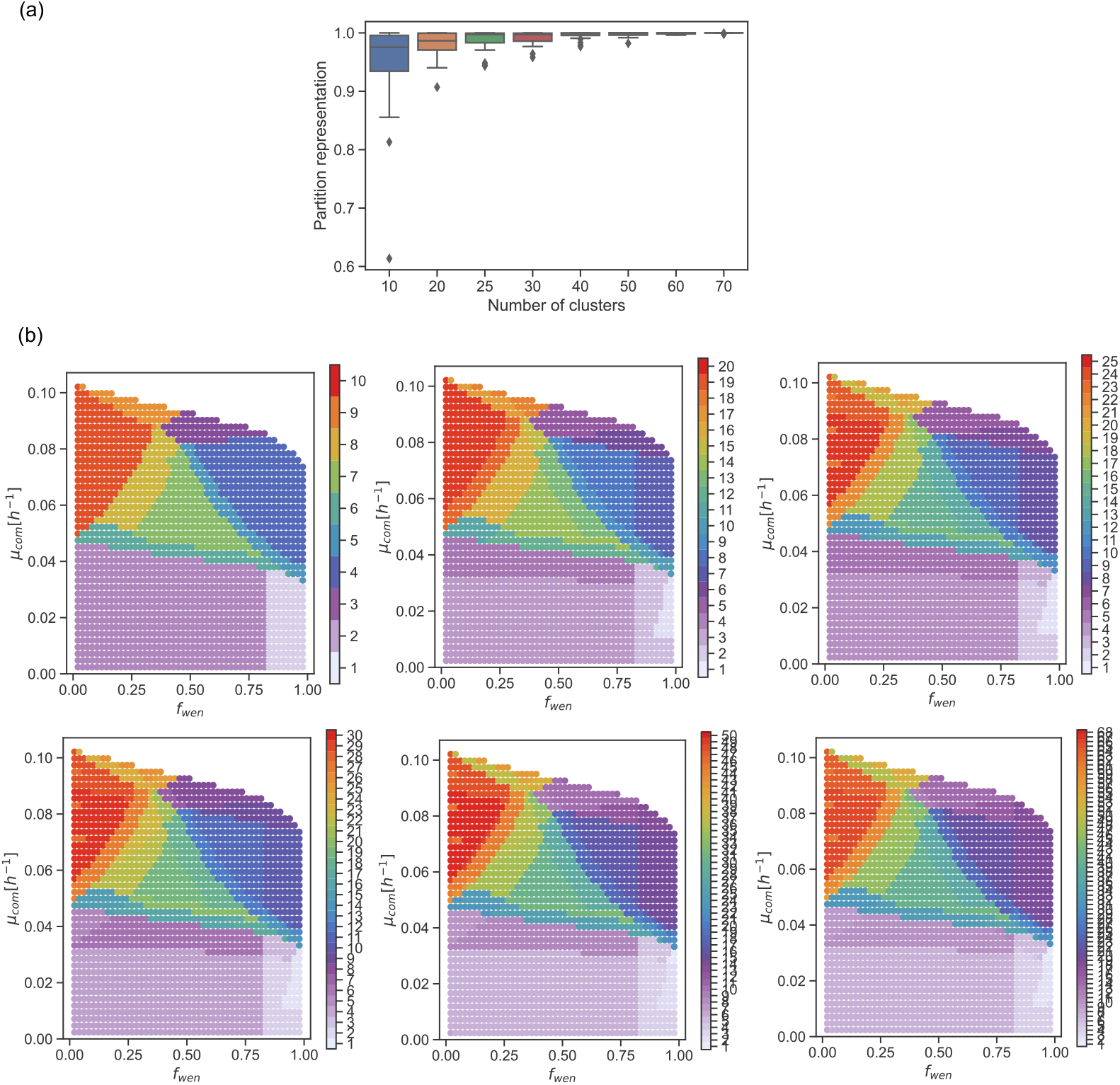
Effect of the variation of the number of clusters. (a) Partition representation improves by increasing cluster number. At *k* = 20, all reactions are accurately represented in at least 90% points of the grid. (b) Zones are obtained in the abundance-growth space when the number of clusters is moved from *k* = 10 to *k* = 69, equivalent to a partition of this space. *k* = 20 captures most of the qualitative shifts of the abundance-growth space.

**FIG S2.**
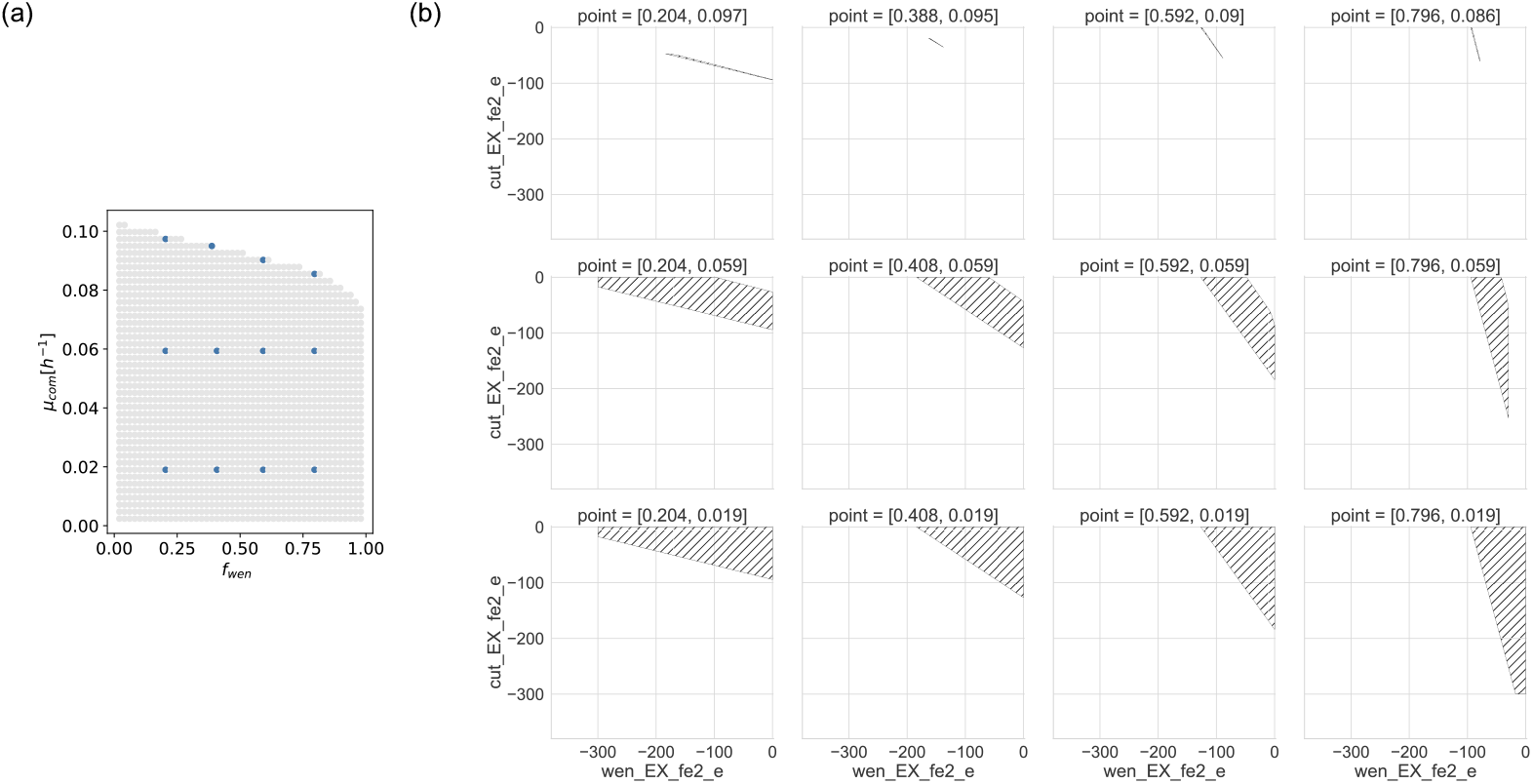
Flux relation analysis - Iron(II) competition. Changes in the feasible fluxes of Fe(II) exchanges (EX_fe2_e) for *A. ferrooxidans* (wen) and *S. thermosulfidooxidans* (cut) computed for specific points in the abundance-growth space. Fluxes are in units of [mmol/gDW_*organism*_ h^−1^]

**FIG S3.**
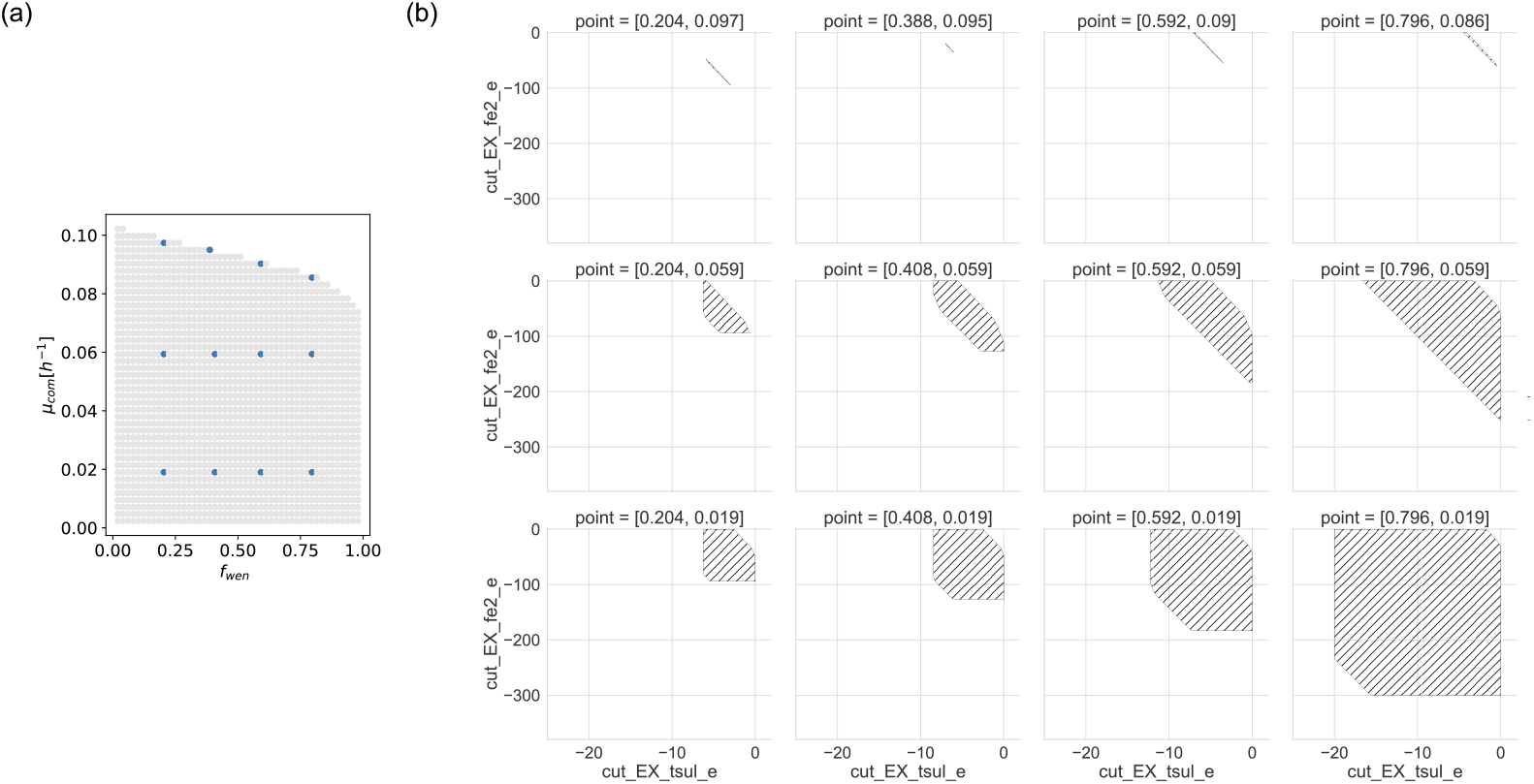
Flux relation analysis - Distribution of resources by *S. thermosulfidooxidans*. Changes in the feasible fluxes of Fe(II) and thiosulfate exchanges (EX_fe2_e and EX_tsul_e) for *S. thermosulfidooxidans* computed for specific points in the abundance-growth space. Fluxes are in units of [mmol/gDW_*cut*_ h^−1^].

**FIG S4.**
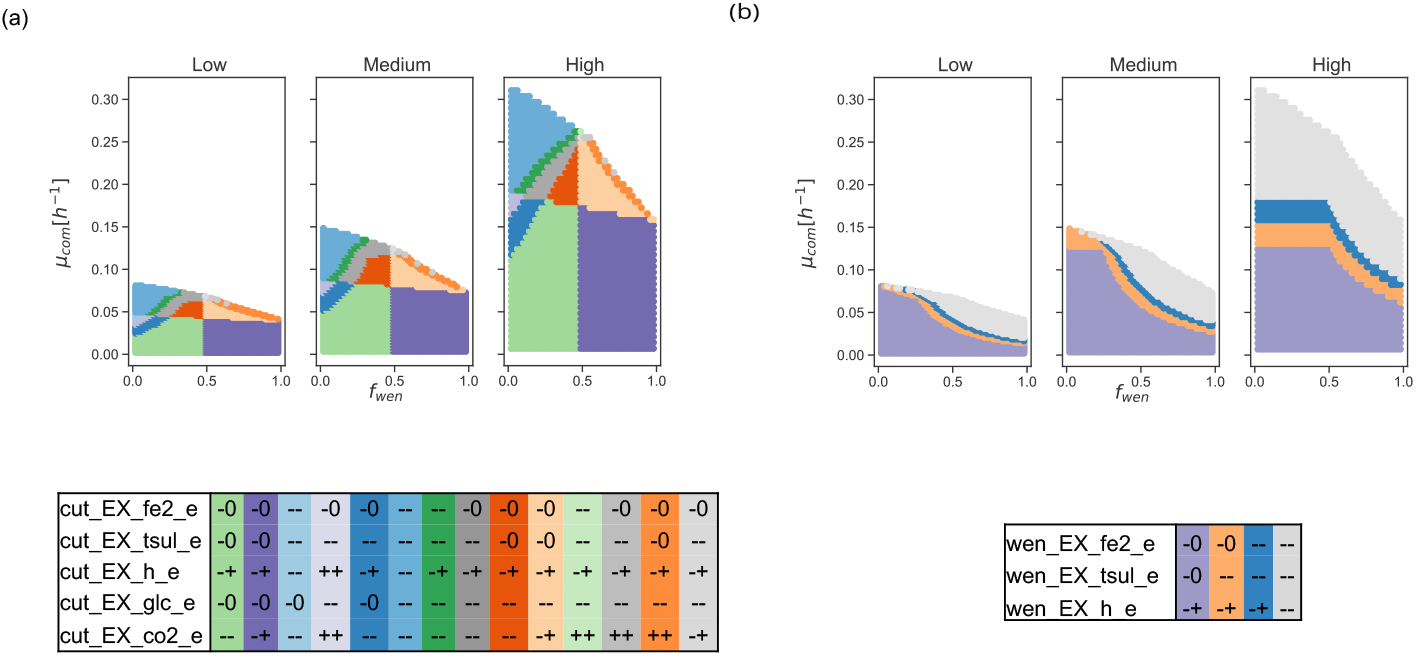
Effect of substrate availability in a high organic availability scenario. A partition of the abundance-growth space defined by qualitative states of key reactions of *S. thermosulfidooxidans* (a) and *A. ferrooxidans* (b) is presented in a scenario with changes in their substrate availability in the presence of high organic matter *α* = 0.6.

FILE S1. Model curation for *A. ferrooxidans* and *S. thermosulfidooxidans*.

FILE S2. Models for *A. ferrooxidans* and *S. thermosulfidooxidans*.

## ACKNOWLEDGMENTS

This workwas supported by the Centerfor Mathematical Modeling(CMM)grant FB210005, which is a Basal fund for centers of excellence from ANID-Chile, Center for Genome Regulation, which is the Millennium Institute Project ICN2021 044 supported by the Millennium Scientific Initiative of the Ministry of Economy, Development and Tourism (Chile), and the National laboratory for high performance computing NLHPC at CMM.

